# Detecting differential growth of microbial populations with Gaussian process regression

**DOI:** 10.1101/055186

**Authors:** Peter D Tonner, Cynthia L Darnell, Barbara E Engelhardt, Amy K Schmid

**Affiliations:** Program in Computational Biology and Bioinformatics, Duke University, Durham, NC 27708, USA; Biology Department, Duke University, Durham, NC 27708, USA; Computer Science Department, Center for Statistics and Machine Learning, Princeton University, Princeton, NJ 08544, USA

**Keywords:** Microbial growth, *Archaea*, machine learning, systems biology, Gaussian processes

## Abstract

Microbial growth curves are used to study differential effects of media, genetics, and stress on microbial population growth. Consequently, many modeling frameworks exist to capture microbial population growth measurements. However, current models are designed to quantify growth under conditions that produce a specific functional form. Extensions to these models are required to quantify the effects of perturbations, which often exhibit non-standard growth curves. Rather than fix expected functional forms of different experimental perturbations, we developed a general and robust model of microbial population growth curves using Gaussian process (GP) regression. GP regression modeling of high resolution time-series growth data enables accurate quantification of population growth, and can be extended to identify differential growth phenotypes due to genetic background or stress. Additionally, confounding effects due to experimental variation can be controlled explicitly. Our framework substantially outperforms commonly used microbial population growth models, particularly when modeling growth data from environmentally stressed populations. We apply the GP growth model to a collection of growth measurements for seven transcription factor knockout strains of a model archaeal organism, *Halobacterium salinarum*. Using these models fitted to growth data, two statistical tests were developed to quantify the differential effects of genetic and environmental perturbations on microbial growth. These statistical tests accurately identify known regulators and implicate novel regulators of growth under standard and stress conditions. Furthermore, the fitted GP regression models are interpretable, recapitulating biological knowledge of growth response while providing new insights into the relevant parameters affecting microbial population growth.

## Introduction

Quantification and prediction of microbial growth is a central challenge relevant to industrial production of value-added chemicals (Lewis et al. 2012), food safety (McKellar and Lu 2003; Ross and Dalgaard 2003), and microbe-environment interactions (Nichols et al. 2011). Parametric models of microbial population growth assume a sigmoid growth function with three characteristic growth phases, captured by three parameters: *lag phase time* (lag phase; λ), during which no growth occurs; *maximum growth rate during logarithmic growth* (log phase; *μ*_*max*_), a phase of rapid growth; and *asymptotic carrying capacity* (stationary phase; *A*), reached when nutrients are exhausted in stationary phase (Baranyi and Roberts 1995; Egli 2009; Monod 1949; Zwietering et al. 1990). Another useful quantification of growth is the area under the growth curve (AUC), also known as *growth potential* (Todor et al. 2014).

Microbial populations frequently encounter shifts away from optimum growth conditions in their environment that require adaptation in order to survive. These shifts, generally referred to as *stress conditions*, include reactive oxygen species (ROS) accumulation, temperature variation, and osmotic fluctuation. These conditions chemically damage or denature macromolecules such as proteins, nucleic acids, and lipids, compromising cellular viability (Imlay 2003; Kühn and Klipp 2012; Verghese et al. 2012). During stress response, the cell state changes from a growth-centric to a survival-centric configuration in which the transcriptional and translational programs are redirected to regulate alternative pathways that repair damage and restore homeostasis (Lu et al. 2009). When stress is severe, or the regulatory program is impaired by mutation, the repair program becomes overwhelmed. In this case, the population growth rate observed by optical density decreases, plateaus, and may even become negative upon cell lysis.

Existing computational methods used to model and predict microbial population growth from time series measurements are parametric functions known as *primary* or *secondary models* (McKellar and Lu 22 2003; Peleg and Corradini 2011; Ross and Dalgaard 2003). Primary models, used to fit data from a population growing on a single main nutrient source (*e.g.*, sugar carbon source), assume the sigmoid growth function. These modeling assumptions lead to inaccurate fits for growth data from cultures exposed to stress conditions that do not have a characteristic sigmoid growth function (Palacios et al. 2014; Sekse et al. 2012). Secondary models were developed to incorporate additional parameters affecting growth, allowing the model to capture stress effects (Peleg and Corradini 2011). The significance of the differences across stress effects under varying growth conditions can be quantified through statistical hypothesis testing (Gommers et al. 1988). However, incorporating these parameters appropriately into these parametric models requires *a priori* knowledge and additional data for how stress perturbations affect growth. For example, a common assumption is that population growth rate follows an Arrhenius equation in response to temperature changes (Barsa et al. 2012). As an alternative to parameteric models of population growth, non-parametric models have been developed to address microbial growth modeling (Cao et al. 2010; di Sciascio and Amicarelli 2008; Palacios et al. 2014); however, many of these models still depend upon parametric primary models in the first stage, an array of parameters based on knowledge of the underlying biological response to growth perturbations, or complicated fitting procedures of the non-parametric model (*e.g.*, optimization of neural net weights). Current models of microbial growth are therefore limited in their general application to novel microbial growth phenotypes.

Across all three domains of life, general stress response mechanisms functioning at the level of gene transcription have been identified that regulate cellular protection and repair (Bonneau et al. 2007; Fiebig et al. 2015; Gasch et al. 2000). These global regulatory programs are induced in response to multiple conditions, and protect cells exposed to one type of stress against others (Jenkins et al. 1988; Lu et al. 2009). Conversely, cells also induce stress-specific responses to aid survival under a particular condition (Stephen et al. 1995; Zuber 2009). The hypersaline-adapted, or halophilic, archaeon *Halobacterium salinarum* is a model organism uniquely suited to study microbial stress response because it survives extremely high levels of UV, ROS, heat shock, and other stressors in its desert salt lake niche (Ng et al. 47 2000; Oren 2008). As such, *H. salinarum* has been extensively studied as a model system for transcription regulatory network architecture and function in response to stress (Schmid et al. 2011, 2009; Todor et al. 2013; Tonner et al. 2015; Todor et al. 2014). A global gene regulatory network computationally inferred from transcriptomic data predicts that over 70 transcription factors (TFs) may control genes whose products adjust physiology and repair damage incurred by stress (Bonneau et al. 2004; Brooks et al. 2014). Follow-up studies that empirically test network predictions have characterized the full set of TF target genes (the “regulon”) and physiological roles of transcription factors that control the response to conditions including oxidative stress through RosR and AsnC (Plaisier et al. 2014; Sharma et al. 2012; Tonner et al. 2015), nutrient availability through TrmB (Schmid et al. 2009; Todor et al. 2014, 2013), metals through SirR (Kaur et al. 2006), iron homeostasis through Idr1 and Idr2 (Schmid et al. 2011), and copper response through VNG1179C (Plaisier et al. 2014). Despite this knowledge, the cellular regulators of growth that respond to environmental perturbation remain understudied in *H. salinarum* and other *Archaea*. For example, the phenotypic impact of mutations to known transcription factors under alternate stress conditions—and the downstream effect of those mutations on the function of the global regulatory network—remain unclear for *H. salinarum* (Brooks et al. 2014) and many other understudied microorganisms (Yoon et al. 2011; Yoon et al. 2013).

Here, we develop a general Gaussian process (GP) regression model of microbial growth to overcome the limitations of parametric growth modeling and to test for differential growth phenotypes of TF mutants under standard and stress conditions. Gaussian processes (GPs) are distributions on arbitrary functions, where any finite number of observations of the function are distributed as a multivariate normal (MVN) in a computationally tractable framework (Rasmussen and Williams 2006). Because GP regression fits an arbitrary functional form, it is able to model growth curves that deviate from the primary sigmoid form. We establish the ability of GP regression to accurately model growth curves from *H. salinarum* under standard and stress treatments across genetic backgrounds. We compare our model with several primary parametric models such as Gompertz, Schnute, and Richards, among others (Zwietering et al. 1990), for growth curve fitting. Subsequent analysis of the GP model output recovers biologically interpretable metrics of microbial growth, including growth rate and carrying capacity. We developed a statistical test of differential growth response between two experiments *via* data likelihoods computed from the fitted GP regression model. We call this model and associated testing framework Bayesian Growth Rate Effect Analysis and Test (B-GREAT). B-GREAT recapitulates known phenotypes of knockout mutants in *H. salinarum* and identifies novel phenotypes, including implicating a metal-responsive transcription factor in oxidative stress resistance.

## Results

We developed a Gaussian process (GP) regression model to capture population growth data from seven *H. salinarum* transcription factor (TF) mutants (Table 1). The growth of these strains was compared to the Δ*ura3* parent strain (from which these mutants were derived) under optimum nutrient conditions (referred to as “standard conditions”) and chronic oxidative stress (see Materials and Methods). Optical density (OD), which quantifies cell density, was measured using a high throughput plate reader (Fig. 1, Supplemental Fig. S1). Population growth phenotypes were measured in a minimum of twelve samples per mutant per condition, sampled every thirty minutes over forty-eight hours for a total of 12,720 data points (Supplemental Table S1). Chronic oxidative stress was induced by the addition of 0.333 mM paraquat (PQ) when the culture was inoculated. The growth rate of these TF strains under standard conditions during log phase has been tested previously (Kaur et al. 2006; Plaisier et al. 2014; Schmid et al. 2009; Schmid et al. 2011; Sharma et al. 2012), but only the growth rates of TF knockout mutants Δ*asnC*, Δ*trmB*, and Δ*rosR* have been tested under PQ conditions (Plaisier et al. 2014; Sharma et al.2012; Table 1). These strains were chosen because the prior studies allow us to validate our results on these previously characterized strains and to discover novel phenotypes.

**Table 1.** **Strains used in this study and their previously known phenotypes and functions**. All phenotypes were previously quantified only in log phase.

| Strain name | Genotype | Phenotype, condition | Pathways regulated | Reference |
| --- | --- | --- | --- | --- |
| $\Delta ura3$ | Parent strain | Control | Control | (Peck et al. 2000) |
| $\Delta trmB$ | $\Delta ura3 \Delta trmB$ | Slow growth, standard<br>Same as control, PQ | Metabolism | (Schmid et al. 2009;<br>Todor et al. 2014;<br>Todor et al. 2013) |
| $\Delta rosR$ | $\Delta ura3 \Delta rosR$ | Slow growth, standard<br>Slow growth, PQ | Oxidative stress repair | (Sharma et al. 2012;<br>Tonner et al. 2015) |
| $\Delta idr1$ | $\Delta ura3 \Delta idr1$ | Same as control, standard | Iron homeostasis | (Schmid et al. 2011) |
| $\Delta idr2$ | $\Delta ura3 \Delta idr2$ | Same as control, standard | Iron homeostasis | (Schmid et al. 2011) |
| $\Delta sirR$ | $\Delta ura3 \Delta sirR$ | Same as control, standard | Manganese uptake | (Kaur et al. 2006) |
| $\Delta VNG1179C$ | $\Delta ura3 \Delta VNG1179C$ | Same as control, standard | Copper uptake | (Kaur et al. 2006) |
| $\Delta asnC$ | $\Delta ura3 \Delta asnC$ | Slow growth, standard<br>Slow growth, PQ | Oxidative stress repair | (Plaisier et al. 2014) |

## Gaussian process regression model of microbial population growth

In order to model the diverse phenotypes observed under both standard and oxidative stress conditions, a probabilistic model of population growth was constructed using Gaussian process (GP) regression (Fig. 1, Supplemental Fig. S1). GP regression is a Bayesian non-parametric model that describes the distribution over an infinite dimensional function *f*(*x*), of which any finite number of observations have a MVN distribution (see Materials and Methods) (Rasmussen and Williams 2006). The GP model is described by its prior mean and covariance functions (*μ*(*x*) and *κ*(*x*, *x*′), respectively). In this study, *μ*(*x*) was set to 0, as is standard (Rasmussen and Williams 2006). The radial basis function (RBF), *κ*(*x*, *x*′), defining the covariance matrix of this MVN distribution, was used as the kernel function. The length scale parameter, *l*, of the RBF kernel specifies the rate of the exponential decay on the covariance between two points *x* and *x*′ (see Materials and Methods). Through the covariance function, GP regression places a prior on all arbitrary functions mapping time to optical density, with functions that have a covariance reflecting the kernel parameters having a high posterior probability. In addition to the covariance defined by the kernel function, independent and identically distributed (IID) Gaussian noise with mean 0 and variance σ^2^ is added to each observation 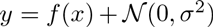. Estimating parameters of a GP regression model on microbial growth data was performed by maximizing the data likelihood with respect to the kernel function parameters (Rasmussen and Williams 2006). We refer to our model (and associated tests, described below) as Bayesian Growth Rate Effect Analysis and Test (B-GREAT).

## GP regression outperforms primary growth models

B-GREAT was used to fit time series growth data from *H. salinarum* Δ*ura3* parent strain populations under both standard and oxidative stress conditions. In order to benchmark GP regression as a model of microbial population growth, GP prediction error was compared to those from four primary growth models: Gompertz (Zwietering et al. 1990), population logistic regression (Zwietering et al. 1990), Schnute (Schnute 1981), and Richards (Richards 1959) (see Materials and Methods). All of the primary growth models depend on the parameters λ and *μ*_*max*_, corresponding to lag time and maximum growth rate, respectively Baranyi and Roberts 1995; Zwietering et al. 1990), of a sigmoidal growth curve. Gompertz, logistic regression, and Richards models also include a parameter for carrying capacity (*A*). Both the Richards and Schnute models include other parameters that modify the sigmoidal shape of the growth curve but do not have direct biological interpretations — *v* for the Richards model, *a* and *b* for the Schnute model (Zwietering et al. 1990).

In order to test model accuracy of GP regression against primary growth models, data were split into training and testing sets comprising 80% and 20% of the data, respectively. Cross validation error, calculated using mean squared error (MSE) between testing data and model prediction based on training data, was calculated for each model under both standard conditions and oxidative stress. As expected, the fit to the data from all models was qualitatively (Fig. 1A) and quantitatively (according to MSE, Fig. 1B) accurate under standard conditions. However, chronic oxidative stress modified the growth trajectory of *H. salinarum* populations such that model predictions deviated from the data (Fig. 1C) and MSE increased by a factor of ten (Fig. 1D) across related models. GP regression outperformed primary models in terms of model fit under both standard and stress conditions (Fig. 1 B and D). GP regression MSE under both conditions was significantly lower than MSE for each of the primary models considered, as determined by a one-sided t-test (Table 2). Unlike primary models, GP regression maintained a similar MSE across standard and stress conditions. Primary models performed poorly under stress conditions, as they are built under the assumption of sinusoidal growth curves. Because GP regression does not assume any specific functional form, our GP models growth data from populations grown under standard and stress conditions with equal accuracy.

**Table 2.** Comparison of GP regression mean squared error (MSE) under standard growth (center column) and oxidative stress (right column) to four primary growth models. Values indicate p-value score of a one-sided t-test between MSE of GP regression and each model for each condition.

| Model | Standard growth | Oxidative stress |
| --- | --- | --- |
| Gompertz | $2.33 \times 10^{-7}$ | $4.51 \times 10^{-77}$ |
| Logistic | $3.73 \times 10^{-30}$ | $2.53 \times 10^{-7}$ |
| Richards | $1.10 \times 10^{-8}$ | $1.12 \times 10^{-67}$ |
| Schnute | $2.22 \times 10^{-5}$ | $5.98 \times 10^{-81}$ |

## GP regression recovers parameters of primary growth models

To enable a biological interpretation of GP growth curves and a quantitative comparison with primary parametric model output, growth parameters of primary models, *A*, *μ*_max_, and AUC, were extracted from our fitted GP models (see Materials and Methods). GP estimates of these parameters under standard growth conditions for the Δ*ura3* parent strain were strongly correlated with those from Gompertz regression (0.903 for *μ_max_* and 0.947 for *A*, *p* ≤ 10^−5^ in both cases as determined by Pearson correlation; Fig. 2A and B, respectively). Estimates of A from Gompertz regression were slightly higher than those from GP regression for a subset of samples (Fig. 2B, Supplemental Fig. S2).

GP regression was then used to estimate A, *μ_max_* and AUC for the Δ*ura3* parent strain, and these estimates were compared with parameter estimates for seven TF deletion strains under both standard and oxidative stress conditions. For each strain, we estimated these three growth parameters from the posterior probability of the fitted GP model (Fig. 2B). This analysis was not performed with primary growth models because GP provides more accurate fits under stress conditions (Fig. 1). According to these parameters, some mutant strains differed from the Δ*ura3* parent under standard conditions, while others differed under oxidative stress. For example, *μ_max_* for the Δ*trmB* strain, a known nutrient responsive regulator, was lower than *μ_max_* for the Δ*ura3* strain under standard conditions as expected from previous studies (Schmid et al. 2009; Todor et al. 2013; Todor et al. 2014). Estimates of A and AUC for the Δ*rosR* strain were lower than *A* and AUC for the other strains. These results demonstrate that growth parameters estimated from GP models are biologically relevant and comparable to those estimated using primary models under standard conditions. GP has the added benefit of estimating these parameters accurately for stress conditions, although the biological interpretation may differ from parameters estimated for standard conditions.

**Figure 1.**
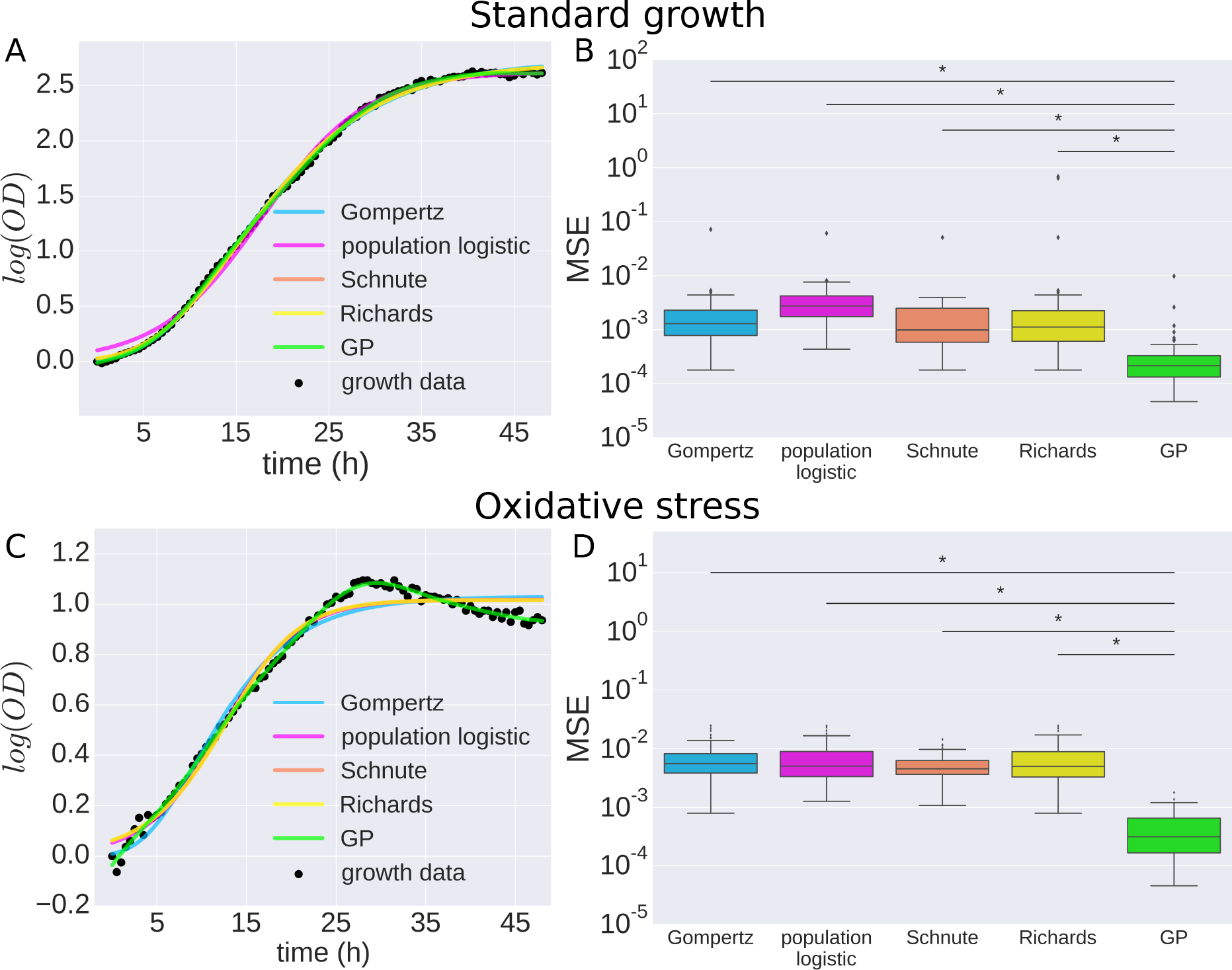
GP regression outperforms primary growth models and recovers parameters of interest. (A) Comparison of GP regression and primary growth models (Gompertz, population logistic, Schnute, Richards) on microbial growth data under standard conditions. (B) Logarithm of mean squared error (MSE) for primary growth models compared to GP regression on microbial population growth under standard conditions. Bars with an asterisk indicate a significant difference between GP MSE and primary growth model MSE as determined by a one-sided t-test. (C) Comparison of GP regression and primary growth models on microbial growth data under oxidative stress. (D) Logarithm of MSE for primary growth models compared to GP regression on microbial population growth under oxidative stress. Bars with asterisks as in (B).

**Figure 2.**
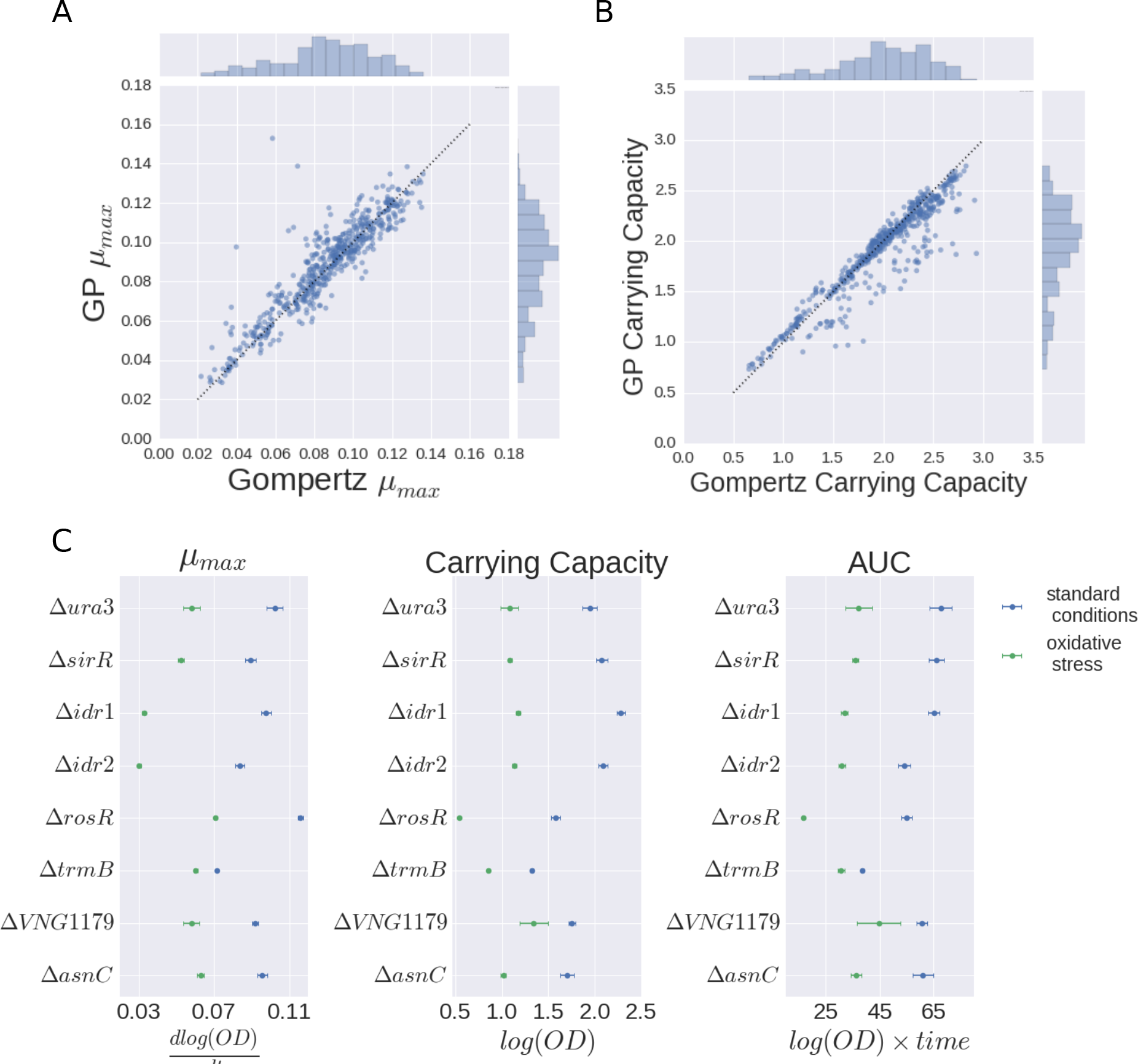
Parameters determined by GP regression. (A) Correlation of parameter estimates of *μ*_*max*_ and (B) carrying capacity (parameter A) between Gompertz and GP regression. Dotted line represents the line *y* = *x*. (C) Posterior distributions of growth parameters *μ*_*max*_, carrying capacity, and AUC are shown for each strain under standard conditions (blue) and oxidative stress (green). Points represent posterior mean function, and error bars indicate 95% credible regions.

## B-GREAT identifies known and novel differential growth phenotypes under 162 standard conditions

Given the superior performance of GP regression to model growth curves, we next sought to identify significant differential growth phenotypes of TF mutants versus the Δ*ura3* parent strain under standard conditions. We developed a statistical test using approximate Bayes factors (BFs) based on our GP regression model. This test was motivated by the observation that, although the growth parameters (*μ*_max_, *A*, AUC) estimated from GP regression are useful in qualitative comparisons of strain growth behavior, the different parameters provide conflicting information in some cases. For example, consider the parameter estimates of Δ*rosR*, where **μ*_max_* estimates were higher than those of the Δ*ura3* parent, while estimates of *A* show growth impairment compared to the Δ*ura3* parent (Fig. 2C). As such, determining whether differential growth is statistically significant can depend on which parameters are examined.

In contrast, our approximate BF test was designed to compare growth curves across specific covariates in the GP regression modeling framework, to capture differences across the entire time series. Specifically, the BF compares the data likelihood under two models, the null and alternative models. We compute approximate BFs by considering point estimates of the GP regression parameters instead of integrating over their uncertainty for computational efficiency. For a null model, we used *f* (time). For an alternative model, we used *OD*(time, strain) = *f* (time, strain), which represents the function of the optical density at a given time and for a specific strain, where a strain value of 0 or 1 indicates parent strain or mutant strain, respectively. The covariate *strain* was added to the model by extending the RBF kernel of the GP to an additional input dimension (Rasmussen and Williams 2006). In this experiment, the null model assumes that the population growth under the condition of interest is the same between parent and mutant strain, while the alternative model assumes that a given mutant population has a different growth response phenotype than the parent strain. Typically, a BF greater than one indicates evidence for the alternative hypothesis, indicating differential growth across the covariate (Kass and Raftery 1995).

In order to compute the statistical significance for our test for differential growth, we used permutations to calibrate the false discovery rate (FDR) of our BFs. To do this, we developed a permutation framework to quantify the distribution of the test statistic under a null hypothesis. We performed calibration *via* permutation in lieu of using a test statistic that has an approximate χ^2^ distribution for more precise calibration at the cost of additional computation (Fusi and Listgarten 2016). Using an estimate of the distribution of the test statistic under the null hypothesis, we quantified the FDR for a given test statistic threshold (Mangravite^*^ et al. 2013). Specifically, across growth data for both parent and mutant strains, the label of strain background was randomly permuted for each time point. Values were permuted so as to maintain the underlying distribution of strain labels present in the original data. 100 permutations were computed for each BF test, and BF scores corresponding to FDR < 20% were considered significant.

**Figure 3.**
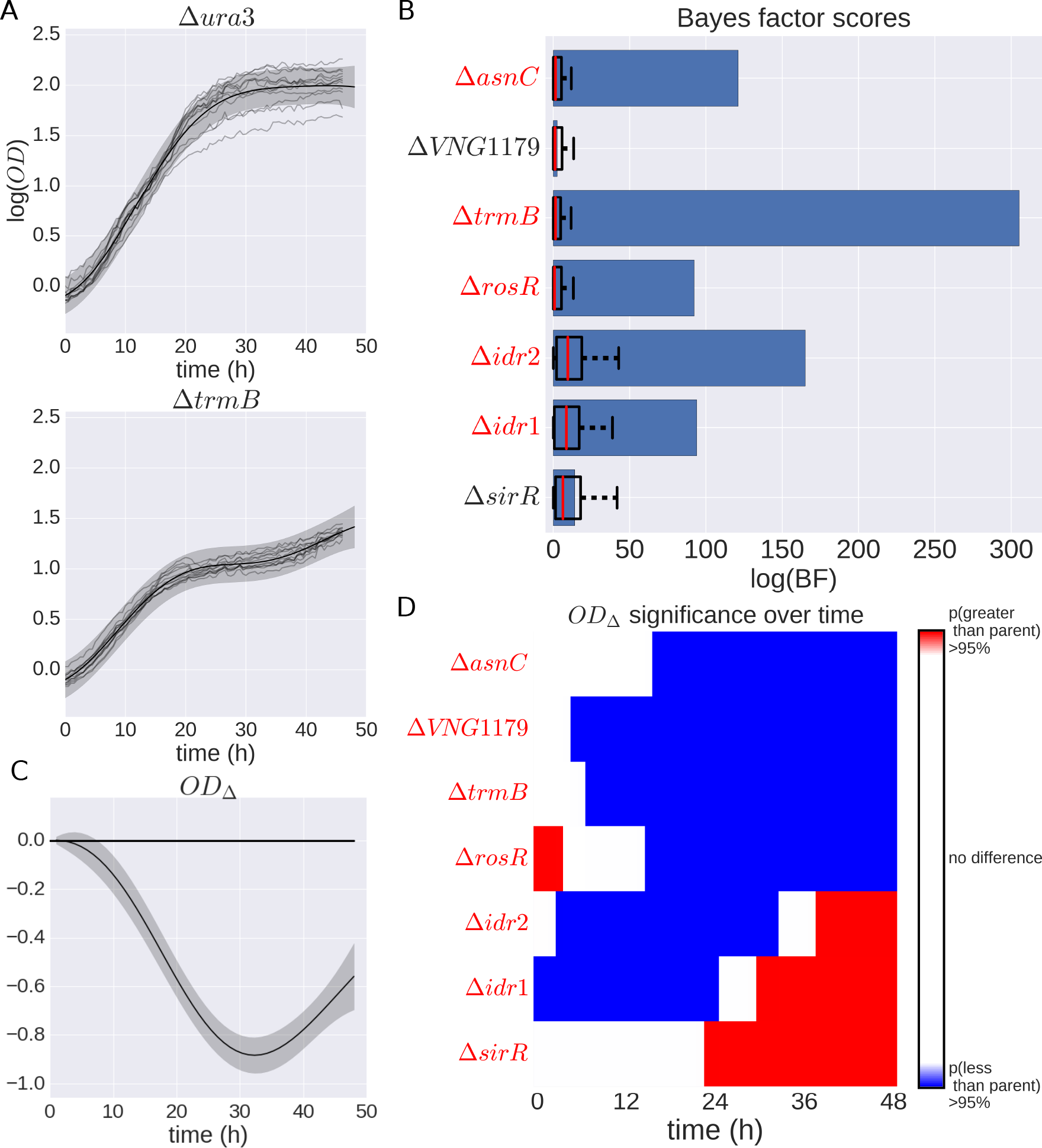
*H. salinarum* mutants with significant growth phenotypes under standard conditions. (A) Population growth data and GP model fit of *H. salinarum* parent strain Δ*ura3* (top) and *ΔtrmB* (bottom) under standard growth. Light grey curves represent growth samples of each strain in different wells. Solid black lines and shaded grey region indicate mean and 95% credible region of the GP model fit to the growth data, respectively. A single GP model was fit (Eq. 2) and separate growth predictions made for Δ*ura3* and Δ*trmB* (see Materials and Methods). (B) Bayes factors for each mutant strain are shown as blue bars. Permuted BF scores are shown as black whisker plots; whiskers represent 1st and 3rd quartiles. Strains with a BF score with FDR ≤ 0.2 are in red italics. (C) The difference in growth level between Δ*trmB* and Δ*ura*3 using the prediction of growth from the GP model. Solid line indicates mean difference and shaded region is the 95% credible region. Regions where the 95% credible region does not include 0 indicate high probability that the growth between the two strains is different at that time point. (D) Predicted difference between mutant and parent strain population growth using posterior function distributions as in (C). Red and blue regions indicate a > 95% probability that the mutant population growth is either higher or lower than the parent strain, respectively. Strains with *OD*_Δ_ 95% credible region not including 0 at any time point are in red italics.

BF scores calculated from GP model fits on growth curves for each mutant strain represent the overall effect of the strain background on population growth (Fig. 3A, B). B-GREAT revealed that five of the seven TF mutants had significant BFs under standard growth conditions, meaning that the mutant strain showed differential growth compared with the parent strain (FDR ≤ 20%), including Δ*asnC*, Δ*trmB*, Δ*rosR*, Δ*idr*2, and Δ*idr*1 (Fig. 3B).

To gain further biological insight into the phenotypes of the five strains with differential growth, we developed a second metric, *OD*_Δ_, that quantifies the difference in parent and mutant strain population growth at each time point of the time course (see Materials and Methods, Eq. 5; Benavoli and Mangili 2015). This difference is computed using the posterior estimates of parent and mutant strain growth of the fitted GP. As we are interested in differences in the actual growth of strain populations, and not differences arising from noise in growth measurements, *OD*_Δ_ is computed using posterior estimates of the underlying growth function without Gaussian noise. The posterior function estimates, and the difference between these estimates, have a MVN distribution. Specifically, we computed the probability of the mutant strain growth conditioning on the parent strain growth at each observation time point according to the MVN distribution. We thresholded this probability at 95% to capture a growth difference between parent and mutant strain at each time point.

As expected from previous work (Schmid et al. 2009), *OD*_Δ_ indicated that Δ*trmB* grows more slowly than the *Aura3* parent strain throughout the time course (Fig. 3C). In contrast, Δidr1 and Δ*idr2* grow more slowly than the parent strain during exponential phase, but reach higher cell densities during the latter portion of the growth curve (Fig. 3D). Δ*rosR* exhibits the opposite growth pattern. The fifth strain with a novel differential growth phenotype, Δ*asnC*, is impaired for growth throughout the time course. Although the growth of Δ*idr*1, Δ*idr*2, Δ*rosR*, and Δ*asnC* strains has been studied during log phase under standard growth conditions previously (Plaisier et al. 2014; Schmid et al. 2011; Sharma et al. 2012), these represent novel stationary phase and stress phenotypes. Taken together, these results demonstrate that B-GREAT and the *OD*_Δ_ metric provide a simple, biologically interpretable test of significance of differential growth that captures the complexity of differential growth phenotypes.

## Identification of differential growth phenotypes in response to oxidative stress

Given that several TF mutants of interest are known to regulate genes in response to stress, such as Δ*rosR* (Sharma et al. 2012; Tonner et al. 2015), we next used B-GREAT to quantify the change in population growth of the TF mutants and Δ*ura3* under chronic oxidative stress. This stressor was introduced to cultures of each strain by adding 0.333 mM paraquat (PQ), a redox cycling drug that permeates the cell membrane. The previous model of growth, *f*(time, strain), was extended to include an effect of PQ and an interaction term between strain and PQ stress: *f* (time, strain, mM PQ, (mM PQ × strain)) (Eq. 3). Here, mM PQ ∈ {0,1} represents the presence or absence of oxidative stress in the culture (Fig. 4A, green curves). Interaction term mM PQ × strain ∈ {0,1} is equal to 1 only for the mutant strain under oxidative stress, and 0 otherwise, and was included to test for differential growth of each mutant strain specific to oxidative stress (Fig. 4A, right blue curve). A BF for this condition was constructed by calculating the relative likelihood of the data with or without the interaction term mM PQ × strain (alternative and null models, respectively). This test statistic quantifies differential strain growth under oxidative stress while controlling for differences in growth between parent and mutant strain under standard conditions. The second test, *OD*_Δ_, was computed as the difference between mutant strain growth with or without the interaction term mM PQ × strain (Fig. 4B).

Using this extended B-GREAT framework, each strain was tested for significant growth phenotypes under PQ stress (Fig. 4C, D). Under this test, Δ*sirR* was the only strain to exhibit a significant difference in growth phenotype under oxidative stress according to *OD*_Δ_ or BF (Fig. 4C, D). According to *OD*_Δ_, Δ*sirR* is impaired for growth relative to the parent strain during the late stages of the growth time series (Fig. 4B). Δ*sirR* was previously implicated in regulating genes involved in metal ion uptake (Kaur et al. 2006), but not in oxidative stress. All other strains were determined to have no significant growth impairment or improvement under PQ stress when differences in strain growth under standard conditions were controlled for in the model (Supplemental Fig. S4). These results indicate that B-GREAT can trivially be extended to include additional covariates, such as strain and interaction term, to enable the discovery of novel growth phenotypes for previously characterized TF mutant strains.

**Figure 4.**
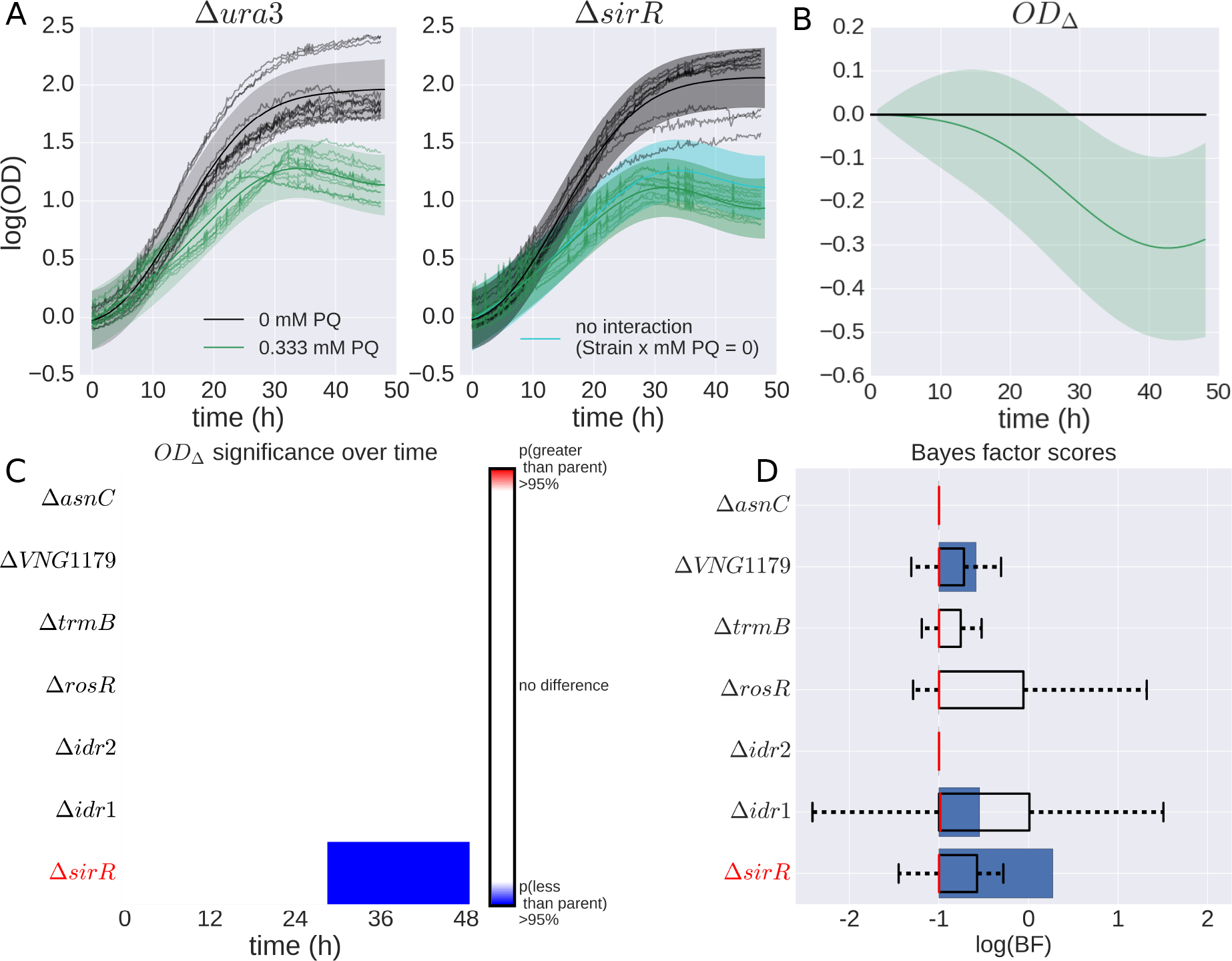
*H. salinarum* mutants with significant growth phenotypes under oxidative stress. (A) Example of population growth data from *H. salinarum* for mutant strain Δ*ura3* (left) and Δ*sirR* (right) under standard conditions (black) and chronic oxidative stress (green). Each curve represents a different sample of an experimental condition. Gaussian process predictions for these conditions are shown as a solid line (mean) and shaded region (variance). The blue line represents the growth prediction when the Strain × mM PQ interaction term is 0. (B) Difference computed between the mutant growth level with interaction term (Strain × mM PQ = 1) and mutant growth without interaction (Strain × mM PQ = 0), solid lines represent mean and shaded regions indicate 95% credible regions. (C) Functional difference and permuted BF scores for mutant strains in response to oxidative stress. Functional difference is computed between mutant strain with and without an interaction term between mutant and stress condition. (D) BF score and permuted BFs for each strain are shown, where bars, boxes, and whiskers are as in Fig. 3C.

## Meta-analysis improves differential growth phenotype detection

Surprisingly, the strain Δ*rosR*, a known oxidative stress regulator that has previously been shown to regulate oxidative stress under both paraquat and hydrogen peroxide exposure (Sharma et al. 2012; Tonner et al. 2015), did not exhibit a significantly differential growth phenotype *versus* the parent strain under oxidative stress (Fig. 4C, D). In order to determine the source of this discrepancy, we compared the growth data for Δ*rosR* generated for this study to that from a previously published study (Supplemental Fig. S5). We observed that Δ*ura3* reached a higher cell density in stationary phase than Δ*rosR* under standard conditions, yielding a significant BF score in our study (Fig. 3B). Thus, controlling for the differential growth under standard conditions removed the differential stress condition phenotype. This difference during stationary phase under standard conditions was observed but not quantified in the previous study because only log phase was considered there (Sharma et al. 2012).

In order to combine the data from this study and from the previous study, we built a hierarchical GP model of growth that corrects for differences arising between batches of experiments (see Materials and Methods) (Hensman et al. 2013). Under this hierarchical model, an underlying growth function *g*(·) is estimated using a GP whose covariates match those in Eq. 3. Then systematic variation between the two studies was modeled as two GPs *f*_1_ and *f*_2_, whose means are given by the shared growth function *g*(·). Under this design, *g* represents the true growth phenotype of Δ*rosR* when corrected for study effects and *f*_1_ and *f_2_* represent the growth phenotype with study-specific effects included (Fig. 5A). From this model, we calculated the difference in Δ*rosR* growth with and without the (mM PQ × strain) interaction term. Once the variation between studies was corrected for, both OD_Δ_ and BF scores (FDR ≤ 0.2) for this model indicate that Δ*rosR* has a significant growth defect under oxidative stress (Fig. 5B, C). This differential phenotype is consistent with the established function of the RosR TF as a genome-wide 270 regulator of gene expression in response to oxidative stress (Tonner et al. 2015). These results demonstrate that this hierarchical model effectively combines cross-study data and corrects for study-specific effects, recapitulating the known phenotype of Δ*rosR* under PQ stress (Fig. 5C).

## Discussion

In this study, we developed a general model of microbial population growth using Gaussian process regres-275 sion to overcome the limitations of commonly used primary parametric models and to enable discovery of novel growth phenotypes for genetically and environmentally perturbed microbial populations. Although primary models provide accurate estimates of growth under optimum culturing conditions where the 3-phase sinusoidal assumption (lag, log, stationary) holds true (Fig. 1A, B), GP accuracy is significantly higher than that of primary models (Fig. 1B). GP regression can recover growth statistics of log phase (*μ*_*max*_) and stationary phase (carrying capacity, A), enabling direct comparison of these variables to results from primary growth models (Fig. 2). Such comparisons revealed that Gompertz regression overestimates *A* for a subset of growth curves, which may contribute to the observed difference in error (Fig. 2B, Supplemental Fig. S2). Rather than extrapolating to unseen data, GP regression advantageously estimates the maximum growth data from the sample provided. In cases where stress conditions change the form of the growth curve and violate the sigmoidal assumption (Fig. 1C), primary models provide a substantially less accurate estimate than that of GP. Taken together, these comparisons demonstrate that GP regression outperforms primary parametric growth models, both under standard conditions and under non-standard stress conditions (Fig. 1).

**Figure 5.**
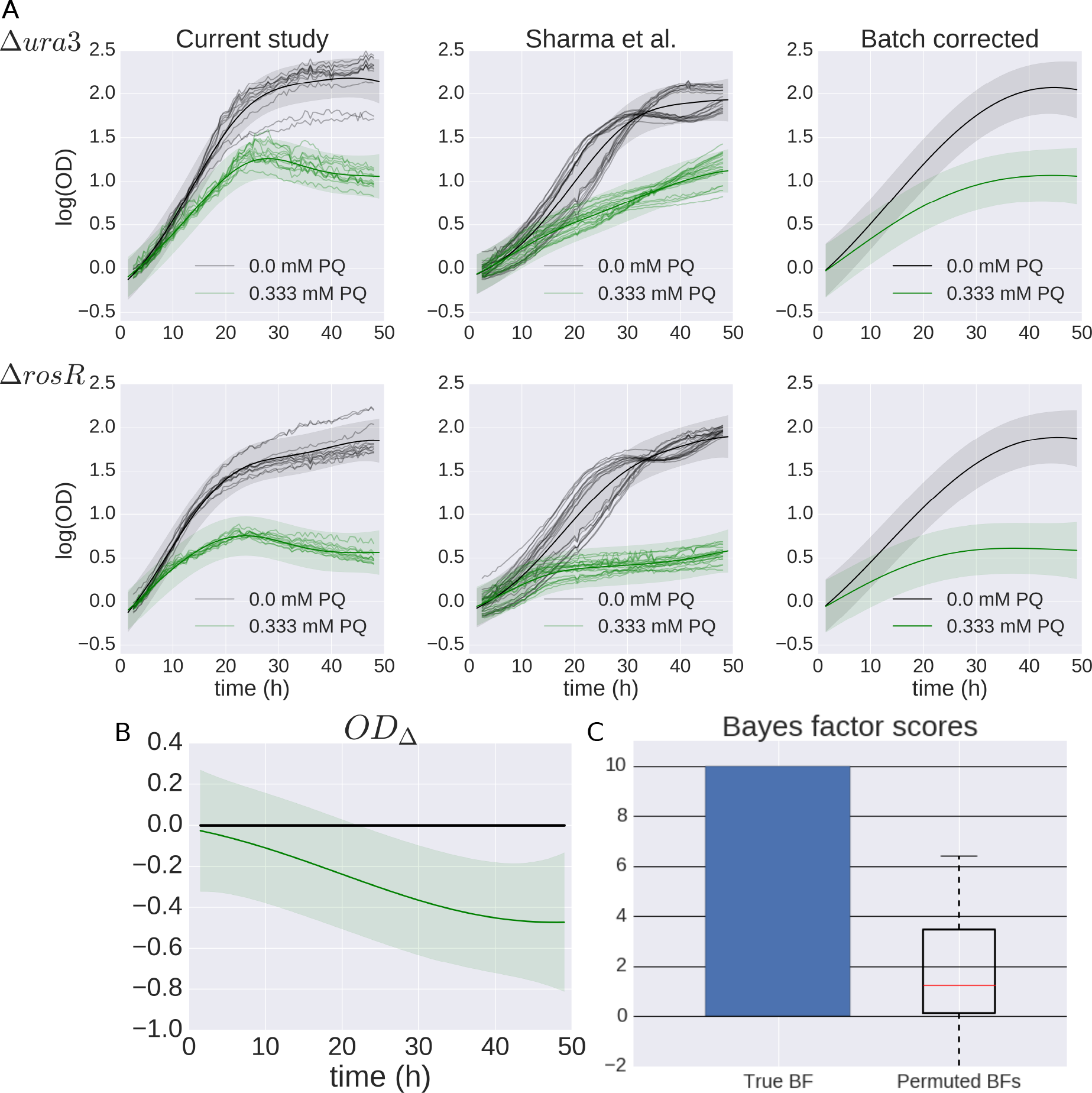
GP model of Δ*rosR* growth in response to oxidative stress across multiple studies. (A) Δ*ura3* (top) and Δ*rosR* (bottom) growth data under standard conditions (black) and oxidative stress (green). Individual samples from this study (left) and previously published data (center, Sharma et al. (2012)) are shown as shaded lines. GP model prediction for each condition is shown as solid line and shaded region for mean and 95% credible region, respectively. The growth prediction for the underlying growth function estimated across studies is shown in the right column. (B) The difference between Δ*rosR* and Δura3 growth for the underlying growth function corrected for batch effects, which shows an increased susceptibility of Δ*rosR* to oxidative stress relative to the parent strain. (C) Bayes factor score compared to permuted scores from the null distribution. Bars, box, and whisker plots as in Fig. 3.

Recovery of growth parameters from GP models is useful for qualitative assessment of growth differences; however, we observed that the magnitude and direction of growth differences can vary across the duration of the time series (*e.g.*, faster or slower growth than wild type; Fig. 2), yielding conflicting results for some parameters. More recently, there has been some attempt to model growth more generally, for example through the use of linear spline regression over portions of the growth curve in the context of a generalized additive model (GAM) (Sekse et al. 2012). Although these models accurately fit growth curves with unexpected functional forms, they are often sensitive to deviations in the growth curve such as those arising from technical variability. Thus, GAM models cannot easily be adapted to testing of differential growth phenotypes.

To overcome these limitations, we developed two metrics for comparing population growth phenotypes, Bayes factors (BFs) and *OD*_Δ_. BFs are a metric in Bayesian analysis for determining the likelihood of two models given data (Kass and Raftery 1995). The *OD*Δ metric provides additional information on the timing and magnitude of growth differences between two conditions. Our use of *OD*Δ is similar to the methods presented in a previous study, in which a hypothesis test was constructed for the equality of two observed functions based on the posterior estimated difference in two GP models fit to each functional observation (Benavoli and Mangili 2015). The previous work concluded that the two functions are the same if the posterior credible region contains the zero vector.

Here, we extended these types of tests, using the posterior difference estimate to quantify the difference between parent and mutant strain growth across the time course, while conducting the hypothesis test for differential growth using approximate BFs. In general, we find that the use of BFs is a conservative method of finding significantly different growth phenotypes relative to *OD*_Δ_, as there are cases in which the posterior difference credible region does not contain zero but the BF is not significant (*e.g.*, Δ*VNG1179C*, Fig. 3). As such, *OD*_Δ_ and BF tests provide two tiers of statistical confidence that an experimental researcher can use to prioritize strains or conditions to pursue for further study. These two metrics are complimentary, since *OD_Δ_* quantifies differenes at each time point, whereas BFs provide a single summary statistic for significant differences across time. Together they capture the complexity of the differential growth phenotype.

With the addition of covariates to our GP model, we tested and accurately identified differential 317 growth across genetic backgrounds (Fig. 3), environmental stress (Fig. 4), and different experimental studies (Fig. 5). With these covariates (Rasmussen and Williams 2006), the GP model learns the independent effects of each perturbation on growth rather than requiring a specification of the parametric effect of perturbations on growth. By adding an interaction term (*e.g.*, strain × stress), the GP model also captures any synergistic effects of genetic background and stress. This interaction is an important consideration given that the number of strains with differential effects increases from one to five out of seven if we remove the mM PQ × strain interaction term (Fig. 4C, Supplemental Fig. S6). Thus, the interaction term is a conservative measure of stress response because it quantifies the impact of strain and stress independently in order to detect significant phenotypes specific to stress while controlling for known differential growth between strains under standard conditions. Given the large proportion of cellular machinery whose production correlates linearly with growth rate (Pedersen et al. 1978; You et al. 2013), differentiating general growth impairments from specific, stress-related impairments is important for biological interpretation of model fits. In terms of correcting for study-specific effects, adding a covariate was not necessary in most cases analyzed here given that other TF mutant strains were collected from a single experimental batch. In contrast, the Δ*rosR* mutant has been analyzed previously (Sharma et al. 2012; Supplemental Fig. S5), and we observed data heterogeneity arising from experimental variation between studies (Fig. 5). To correct for this, we extended the BF using hierarchical GP regression (Hensman et al. 2013), which explicitly controlled for study effects. In the future, GP regression may be extended by adding new covariates to model other growth conditions, such as gradients of chemical stresses (Sekse et al. 2012).

GP regression recapitulates known biology and discovers previously uncharacterized phenotypes. We confirmed the known growth defect for Δ*trmB* under standard conditions (Fig. 3B-D), which results from its function as a master regulator of metabolic pathways (Schmid et al. 2009; Todor et al. 2013; Todor et al. 2014; Todor et al. 2015). We combined data from this study with data from a previous study (Sharma et al. 2012) to confirm the susceptibility of the Δ*rosR* mutant to oxidative damage (Fig. 5). This phenotype is consistent with the ability of RosR to regulate approximately twenty other genes encoding TFs and oxidative repair genes, enabling cellular viability during oxidative stress (Tonner et al. 2015). In contrast to these results with Δ*trmB* and Δ*rosR*, the Δ*asnC* oxidative stress phenotype observed by Plaiser and colleagues (Plaisier et al. 2014) was not recapitulated here, likely because the growth defect of this mutant under standard conditions explains the difference in growth during stress (Fig. 3C, Supplemental Fig. S3). We note that, while the standard condition differential phenotype of Δ*asnC* was observed previously, it was not corrected for in the earlier work, which likely explains the discrepancy (Plaisier et al. 2014). Finally, we identified a previously undiscovered relationship between Δ*sirR* and oxidative damage (Fig. 4). SirR regulates metal uptake transporters at the level of transcription (Kaur et al. 2006), repressing manganese uptake transporters under replete conditions. As a result, expression of genes encoding metal uptake transporters is constitutively high in a Δ*sirR* mutant. Because metal excess leads to oxidative stress through the Fenton reaction (Imlay 2003), this regulatory link between metal homeostasis and oxidative stress is well-established in bacterial and eukaryotic organisms. However, this connection in *Archaea* is only beginning to be appreciated (Zhu et al. 2013).

## Materials and Methods

### H. *salinarum* growth data

Growth of seven transcription factor (TF) mutant strains for *H. salinarum*, each deleted in-frame for a TF-encoding gene, and the isogenic Δ*ura3* parent strain was measured (Table 1). Details regarding construction of these mutants were described in prior work (Kaur et al. 2006; Plaisier et al. 2014; Schmid et al. 2011, 2009; Sharma et al. 2012). Cultures were inoculated into complete medium (CM; 250 NaCl, 20 g/l MgSO4 7H2O, 3 g/l sodium citrate, 2 g/l KCl, 10 g/l peptone), grown to stationary phase, then diluted to OD ~ 0.05 for growth analysis. Optical density (OD) at 600 nm of 200 independent cultures was measured every thirty minutes for 48 hours using a Bioscreen C (Growth Curves USA, Piscataway, NJ). Growth of each strain under each experimental condition was measured in at least biological quadruplicate (from independent colonies) and technical triplicate (independent cultures from the same colony), for a total of twelve replicates. Standard and chronic oxidative stress conditions were tested for all mutants. Standard conditions were defined as 42°C with 225 r.p.m. shaking under ambient light in rich CM medium (Yao and Facciotti 2011). Chronic oxidative stress was induced with 0.333 mM paraquat (PQ) added at the inoculation of the Bioscreen experiment.

Prior to statistical analysis, OD data were log_2_ transformed and scaled by the estimate of starting OD as follows. Data from growth experiments were grouped by their strain and media composition (*e.g*., Δ*ura3*, standard growth). This corresponds to the twelve replicates comprising four biological replicates and three technical replicates. Then OD measurements from the first ten time points within each group were fit with a polynomial regression of degree five. The OD value at time = 0, as estimated by the polynomial regression, was then subtracted from all data points in the group in order to normalize the starting growth levels at zero for all conditions.

### H. *salinarum* data as input to B-GREAT

Input to the GP model corresponds to measurements *Y_t,c,r_* for a given time (1 ≤ *t* ≤ *T*), condition (1 ≤ *c* ≤ *C*), and replicate (1 ≤ *r* ≤ *R*). For standard conditions, time points were taken at 4 hour increments across a 48 hour experiment. This resulted in 12 observations from each replicate. Additionally, growth measurements from both parent strain and each mutant strain were included (*C* = 2). A total of *T* × *R* × *C* = 288 observations were used for training each GP model under standard conditions. For oxidative stress, time points were taken from every 6 hours, for a total of 8 time points for each replicate. The decrease in time samples used in the oxidative stress models was necessary to incorporate the increase in conditions for both standard and oxidative stress growth. Specifically, conditions include growth for both parent and mutant strain under both standard and oxidative stress conditions (*C* = 4). This corresponds to a total of 384 observations for each GP model under oxidative stress. Gaussian process regression of microbial population growth data

## Gaussian process regression of microbial population growth data

Gaussian process (GP) regression is a probability distribution on arbitrary functions mapping *x* to *f*(*x*) (Rasmussen and Williams 2006). When observations of *f*(*x*) are distorted with independent and identi-cally distributed (IID) Gaussian noise, multiple observations of the function are distributed as a multivariate Gaussian

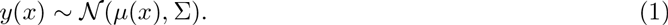

In our application, *x* represents time and *y*(*x*) = *logOD*(*x*) represents the log-transformed OD measurement at time *t*. A GP model requires specification of a mean function *μ*(*x*) and kernel function Σ_*i,j*_ = *κ*(*x*_*i*_, *x*_*j*_), which defines the positive definite covariance matrix Σ. In this work, the mean function was set to zero across inputs, *μ*(*x*) = 0, as is standard (Rasmussen and Williams 2006). For the kernel, we used a radial basis function (RBF) with time point specific independent Gaussian noise:

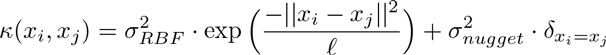

Here, *x_i_* and *x_j_* are two time points, 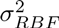 is the RBF variance parameter, 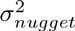 is the Gaussian variance at a single time point *t* (called the *nugget*), 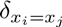 is an indicator function, which is equal to 1 when *x_i_* = *x_i_* = *x_j_* and 0 otherwise, and *l* is the RBF *length scale* parameter, which dictates the smoothness of the function *f*(*x*) through the GP distribution. Kernel function parameters 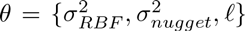 were optimized by maximizing the likelihood of the data marginalized over the latent function *f* (*x*) with respect to each parameter (Rasmussen and Williams 2006). All GP regression models were built and optimized using the GPy package (version 0.8.8) for Python (http://github.com/SheffieldML/GPy).

## GP growth curve metrics

The growth curve metrics *μ_max_* and carrying capacity *A* were calculated from the maximum *a posteriori* (MAP) estimates of either *log*(*OD*) or 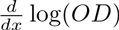 for carrying capacity and *μ_max_*, respectively. MAP estimates of *log*(*OD*) are given by the model in Eq. 1, by taking the MAP growth level using the fitted model. In order to calculate a MAP estimate of 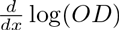, we must estimate 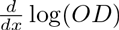 using GP regression. The RBF kernel is infinitely differentiable, so derivative observations of a GP regression model are also distributed as a GP as follows (Solak et al. 2003): 
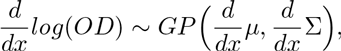
where 
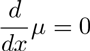
and 
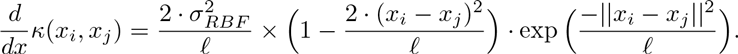

The GP model of 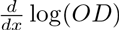 is used to calculate the MAP estimate of 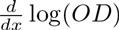 as an estimate of *μ*max.

## Primary growth models

We compared the predictions from the fitted GP regression model to predictions from four primary growth curve models: Gompertz, population logistic, Schnute, and Richards regression (Zwietering et al. 1990). All model parameters were optimized with the `curve_fit` function of the `scipy` Python package, which estimates function parameters using damped least squares (Millman and Aivazis 2011). Data were randomly divided into training (80%) and testing (20%) sets. The mean squared error (MSE) of each model fit with respect to the 20% held out testing data was calculated as the difference between prediction and test data from models built from the training data: 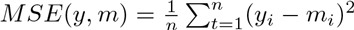, where *y_i_* and *m_i_* correspond to raw data and model predictions at the ith time point, respectively. Model prediction from GP was taken as the posterior mean of the fitted GP, and primary growth model predictions were taken from the growth level predicted by the estimated parameters. Using a one-sided sample t-test, MSE for GP regression fit was compared separately to each of Gompertz, population logistic, Schnute, and Richards regression fits. These primary models were selected to compare against the most widely used primary models in modeling microbial population growth. Additionally, the models chosen have been shown to be related to one another through specific constraints on parameters. For example, Gompertz regression can be recovered from the Schnute model with parameters *a >*0 and *b* = 0 (Zwietering et al. 1990). Therefore, we can observe the improvement of primary model accuracy as we add additional 421 parameters. Gompertz regression.where *A* is the carrying capacity, *μ_max_* is the maximum growth rate, and λ is lag time (Zwietering et al. 1990).Therefore, we can observe the improvement of primary model accuracy as we add additional parameters.

## Gompertz regression

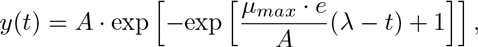
where *A* is the carrying capacity, *μ*_*max*_ is the maximum growth rate, and *λ* is lag time (Zwietering et al. 1990).

### Population logistic regression

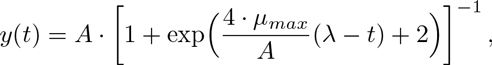
where *A* is the carrying capacity, *μ*_*max*_ is the maximum growth rate, and *λ* is lag time. (Zwietering et al. 1990).

### Schnute model

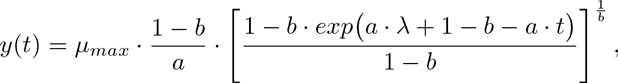
where *μ_max_* is the maximum growth rate, *λ* is lag time, and *a*, and *b* are parameters that affect the growth curve shape but do not have direct biological interpretation (Zwietering et al. 1990).

### Richards model

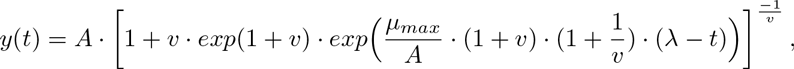
where *A* is the carrying capacity, *μ_max_* is the maximum growth rate, *λ* is lag time, and *v* is a parameter that affects the growth curve shape but does not have direct biological interpretation (Zwietering et al. 1990).

## Testing for differential growth using Bayes factors

We developed an approximate Bayes factor (BF) test statistic to quantify possible differences between a pair of growth conditions BF_strain_ (Kass and Raftery 1995). BFs were calculated as the ratio of data likelihoods between an alternative model (*H*_*a*_) and a null model (*H*_0_):

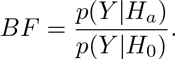

Larger values of the BF indicate a higher relative likelihood under the alternative model and provide evidence for the alternative model representing the data better than the null model.

Specifically, we designed three different BF test statistics to measure differences in population growth across covariates. Under standard conditions, we use BF_strain_, in which the null model *H_0_* assumes that growth is the same across parent and mutant strain; the alternative model *H_a_* captures growth between parent and mutant strain separately. A high BF then suggests that the growth phenotype is different across strains. We designed a second test for differential growth in the presence of oxidative stress, BF_stress_, where the alternative model included an interaction term between genetic effect and oxidative stress. High BF scores under this condition indicate that the mutant strain has a differential growth phenotype relative to the parent strain under oxidative stress. We designed a third test for differential growth across two separate studies, BF_study_, which performs the same test as BF_stress_ but shares statistical strength across batches of growth measurements using a hierarchical GP model.

A false discovery rate (FDR) for each BF was calculated using an estimate of the null BF distribution, representing BF scores when no significant growth effect between the two conditions is observed. For a single growth experiment, *Y* = {*y*_1_,*y*_2_,… *y*_T_}, and corresponding time, genetic background, and other covariates *X* = {*x*_1_, *x*_2_,… *x*_*T*_}, each *x_t_* = {time, strain,…} was randomly assigned a value for *strain* that preserved the original distribution of *strain* values in *X*. 100 permutations of the data indices following this design were constructed, and a BF score was calculated for each permutation. The distribution of permuted BF scores was used as a estimate of the null distribution of the test statistic, and a BF score 451 that exceeded 80% of permuted scores (corresponding to FDR ≤ 0.2) was selected as significant.

More generally, FDR is calculated using permutations, for a given BF threshold *c*, as follows:

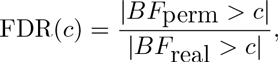

which approximates the FDR, *i.e.*, the number of false positives over the total number of discoveries, for threshold c. In this case, there is a single BF_real_ for 100 permuted BFs, so we multiplied the BF_real_ count by 100 for this computation.

## Differential mutant growth phenotypes

The effects of gene deletion on growth were modeled as experimental effects by extending the input variable *x*, originally representing time, to include perturbations as additional covariates in the GP regression model. The RBF kernel function was extended to handle the additional covariates by using an automatic relevance determination (ARD) prior to induce sparsity on the weighted contribution of each of the *K* covariates (MacKay 1992; Neal 2012; Rasmussen and Williams 2006; Tipping 2001):

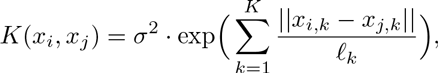

where each *l_k_* is the length-scale for the *k^th^* covariate. These length-scale parameters are then interpretable in terms of quantifying the relative contribution of each of the *k* covariates. Genetic background was incorporated into the model covariates as a Boolean variable x_stran_; ∈ {0,1}, where a value of 0 indicates parent strain and 1 indicates mutant strain. For standard growth conditions, *x* has the form:

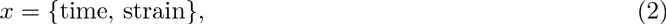

where as the null model contains no strain information: *x* = {time}. The BF then quantified the improvement in data likelihood of the GP regression model including the strain information versus omitting strain information; when modeling strains separately improved the data likelihood, this indicated that there was differential growth across strains.

## Differential response to stress conditions across strains

Differential growth in response to paraquat (PQ) exposure was tested by extending the covariates to include two additional covariates. The first covariate, mM PQ ∈ {0,1}, represents the presence (1) and absence (0) of PQ stress. The second covariate, mM PQ × strain ∈ {0,1} is an interaction term between mutant strain and stress condition, computed by multiplying the strain covariate with the mM PQ covariate. mM PQ × strain covariate was 1 only for growth measurements made under oxidative stress for the mutant strain, and 0 otherwise. The test for significant growth phenotypes was then made using models including or excluding the mM PQ × strain interaction term. Specifically, the input *x* for the paraquat condition had the form:

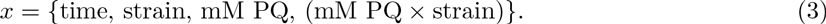

The null model, where there is no interaction between strain and stress condition, corresponds to:

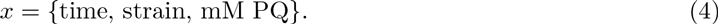

## Modeling batch effects and testing for differential effects across studies

Growth data for Δ*rosR* under standard conditions and oxidative stress was collected both in this study and in a previous study (Sharma et al. 2012). We modeled the joint growth data from both studies with a hierarchical GP model (Hensman et al. 2013). Under this model, the underlying growth function is modeled with a GP: *g*(*x*) ~ *GP*(*μ*_*g*_,*K*_*g*_), and different batch observations of this function are drawn from a GP with mean equal to *g*(*x*): *f*(*x*) ~ *GP*(*g*(*x*),*K*_*f*_).

Growth data for Δ*rosR* and the parent strain were modeled by replicate functions *f*_1_ and *f*_2_, repre-senting data from our study and the previous study, respectively. The GP models for *f*_1_, *f*_2_, and *g* all follow the design in Eq. 3. BF scores in both cases were calculated as the difference in log likelihood for GP models accounting for strain variation interacting with oxidative stress (*HA*; Eq. 3) and those that do not interact with oxidative stress (*H*_0_; Eq. 4). The BF permutation was performed as described above.

## Computing differences between populationc growth across time series (*OD*_Δ_)

The difference between mutant and parent strain functions across time points were defined by the variable *OD*_Δ_. The variable *OD*_Δ_ is the difference in mutant strain growth and parent strain growth at each time point *x*. *OD*_Δ_ was calculated using the noiseless latent mean function for population growth, rather than the noisy observations. In other words, we use the latent function *f*, where *log*(*OD*) = *f*(*x*) + *ϵ* where *f*(*x*) is the smooth underlying growth function and *e* represents random noise. Additionally, *OD*_Δ_ was corrected by the population growth at the start of the experiment, *t*_0_:

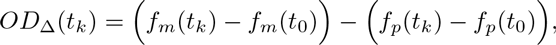

where *t*_0_ denotes the start of the experiment, and *f*_*m*_ and *f_p_* indicate the mutant and parent strain posterior mean function predictions, respectively. The four variables needed to calculate *OD*_Δ_; *i.e.*, *f*_*k*_ = {*f*_*m*_(*t*_*k*_), *f*_*m*_(*t*_0_), *f*_*p*_(*t*_*k*_), and *f*_*p*_(*t*_0_)}; are defined by a joint MVN distribution predicted by the fitted GP:

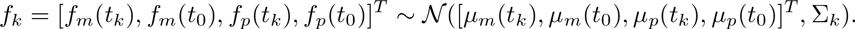

*OD*_Δ_ is then a linear transformation of these variables, *OD*_Δ_ = *a* · *f_k_* where *a* is the column vector *a* = [1, −1, −1,1] (*a*: 1 × 4). Parameter *OD*_Δ_ is then distributed as a univariate normal distribution, 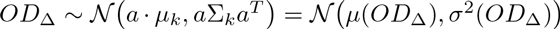. Credible intervals of *OD*_Δ_ as defined by its normal distribution were calculated to determine whether *OD*_Δ_ = 0 lies within the 95% credible region.

## Data access

All code and data associated with this paper are available at https://github.com/ptonner/gp_growth_phenotype. Raw growth data used in this study is available in Supplemental Table S1.

## Acknowledgments

PDT was funded by a National Science Foundation (NSF) Graduate Research Fellowship. BEE was funded by NIH R00 HG006265, NIH R01 MH101822, and a Sloan Faculty Fellowship. AKS was funded by NSF MCB-141-7750. Any opinions, findings, and conclusions or recommendations expressed in this material are those of the author(s) and do not necessarily reflect the views of the National Science Foundation.

## Disclosure declaration

PDT, CLD, BEE, and AKS have no conflicts of interest to declare.

## References

BaranyiJ, Roberts TA. 1995. Mathematics of predictive food microbiology. International Journal of Food Microbiology, 26(2):199–218.

Barsa CS, Normand MD, Peleg M. 2012. On Models of the Temperature Effect on the Rate of Chemical Reactions and Biological Processes in Foods. Food Engineering Reviews, 4(4):191–202.

Benavoli A, Mangili F. 2015. Gaussian Processes for Bayesian hypothesis tests on regression functions. In Conference on Artificial Intelligence and Statistics, volume 38, pp. 74–82.

Bonneau R, Baliga NS, Deutsch EW, Shannon P, Hood L. 2004. Comprehensive de novo structure prediction in a systems-biology context for the archaea Halobacterium sp. NRC-1. Genome Biology, 5(8):1–15.

Bonneau R, Facciotti MT, Reiss DJ, Schmid AK, Pan M, Kaur A, Thorsson V, Shannon P, Johnson MH, Bare JC, et al. 2007. A Predictive Model for Transcriptional Control of Physiology in a Free Living Cell. Cell, 131(7):1354–1365.

Brooks AN, Reiss DJ, Allard A, Wu W, Salvanha DM, Plaisier CL, Chandrasekaran S, Pan M, Kaur A, Baliga NS. 2014. A system-level model for the microbial regulatory genome. Molecular Systems Biology, 10(7):740. DOI:10.15252/msb.20145160.

Cao R, Francisco-Fernndez M, Quinto EJ. 2010. A random effect multiplicative heteroscedastic model for bacterial growth. BMC Bioinformatics, 11:77. DOI: 10.1186/1471-2105-11-77.

di Sciascio F, Amicarelli AN. 2008. Biomass estimation in batch biotechnological processes by Bayesian Gaussian process regression. Computers & Chemical Engineering, 32(12):3264–3273.

Egli T. 2009. Growth Kinetics, Bacterial. In Encyclopedia of Microbiology, 3rd ed. (ed. Schaechter M), pp. 180–193. Academic Press, Oxford.

Fiebig A, Herrou J, Willett J, Crosson S. 2015. General Stress Signaling in the Alphaproteobacteria. Annual Review of Genetics, 49:603–625.

Fusi N, Listgarten J. 2016. Flexible Modelling of Genetic Effects on Function-Valued Traits. In Research in Computational Molecular Biology (ed. Singh M), pp. 95–110. Springer, Switzerland.

Gasch AP, Spellman PT, Kao CM, Carmel-Harel O, Eisen MB, Storz G, Botstein D, Brown PO. 2000. Genomic Expression Programs in the Response of Yeast Cells to Environmental Changes. Molecular Biology of the Cell, 11(12):4241–4257.

Gommers PJF, van Schie BJ, van Dijken JP, Kuenen JG. 1988. Biochemical limits to microbial growth yields: An analysis of mixed substrate utilization. Biotechnology and Bioengineering, 32(1):86–94.

Hensman J, Lawrence ND, Rattray M. 2013. Hierarchical Bayesian modelling of gene expression time series across irregularly sampled replicates and clusters. BMC Bioinformatics, 14(1):1–12.

Imlay JA. 2003. Pathways of oxidative damage. Annual Review of Microbiology, 57:395–418.

Jenkins DE, Schultz JE, Matin A. 1988. Starvation-induced cross protection against heat or H_2_O_2_ challenge in Escherichia coli. Journal of Bacteriology, 170(9):3910–3914.

Kass RE, Raftery AE. 1995. Bayes Factors. Journal of the American Statistical Association, 90(430):773–795.

Kaur A, Pan M, Meislin M, Facciotti MT, El-Gewely R, Baliga NS. 2006. A systems view of haloarchaeal strategies to withstand stress from transition metals. Genome Research, 16(7):841–854.

Kühn C, Klipp E. 2012. Zooming in on Yeast Osmoadaptation. Advances in Experimental Medicine and Biology, 736:293–310.

Lewis NE, Nagarajan H, Palsson BO. 2012. Constraining the metabolic genotype-phenotype relationship using a phylogeny of in silico methods. Nature Reviews Microbiology, 10(4):291–305.

Lu C, Brauer MJ, Botstein D. 2009. Slow growth induces heat-shock resistance in normal and respiratory-deficient yeast. Molecular Biology of the Cell, 20(3):891–903.

MacKay DJC. 1992. Bayesian interpolation. Neural Computation, 4(3):415–447.

Mangravite^*^ LM, Engelhardt^*^ BE, Medina MW, Smith JD, Brown CD, Chasman DI, Mecham BH, Howie B, Shim H, Naidoo D, et al. 2013. A statin-dependent QTL for GATM expression is associated with statin-induced myopathy. Nature, 502(7471):377–80.

McKellar R, Lu X. 2003. Primary Models. In Modeling Microbial Responses in Food, 1st ed. (ed. McKellar R, Lu X), pp. 21–62. CRC Press, Boca Raton.

Millman KJ, Aivazis M. 2011. Python for Scientists and Engineers. Computing in Science & Engineering, 13(2):9–12.

Monod J. 1949. The Growth of Bacterial Cultures. Annual Review of Microbiology, 3(1):371–394.

Neal RM. 2012. Bayesian learning for neural networks, volume 118. Springer Science & Business Media.

Ng WV, Kennedy SP, Mahairas GG, Berquist B, Pan M, Shukla HD, Lasky SR, Baliga NS, Thorsson V, Sbrogna J, et al. 2000. Genome sequence of Halobacterium species NRC-1. Proceedings of the National Academy of Sciences, 97(22):12176–12181.

Nichols RJ, Sen S, Choo YJ, Beltrao P, Zietek M, Chaba R, Lee S, Kazmierczak KM, Lee KJ, Wong A, et al. 2011. Phenotypic landscape of a bacterial cell. Cell, 144(1):143–156.

Oren A. 2008. Microbial life at high salt concentrations: phylogenetic and metabolic diversity. Saline Systems, 4:1–13.

Palacios AP, Marn JM, Quinto EJ, Wiper MP. 2014. Bayesian modeling of bacterial growth for multiple populations. The Annals of Applied Statistics, 8(3):1516–1537.

Peck RF, DasSarma S, Krebs MP. 2000. Homologous gene knockout in the archaeon Halobacterium salinarum with ura3 as a counterselectable marker. Molecular Microbiology, 35(3):667–676.

Pedersen S, Bloch PL, Reeh S, Neidhardt FC. 1978. Patterns of protein synthesis in E. coli: a catalog of the amount of 140 individual proteins at different growth rates. Cell, 14(1):179–190.

Peleg M, Corradini MG. 2011. Microbial growth curves: what the models tell us and what they cannot. Critical Reviews in Food Science and Nutrition, 51(10):917–945.

Plaisier CL, Lo F, Ashworth J, Brooks AN, Beer KD, Kaur A, Pan M, Reiss DJ, Facciotti MT, Baliga NS. 2014. Evolution of context dependent regulation by expansion of feast/famine regulatory proteins. BMC Systems Biology, 8:1–14.

Rasmussen CE, Williams KI. 2006. Gaussian processes for machine learning. In Adaptive computation and, machine learning (ed. Dietterich T). MIT Press, Cambridge, Mass.

Richards FJ. 1959. A Flexible Growth Function for Empirical Use. Journal of Experimental Botany, 10(2):290–301.

Ross T, Dalgaard P. 2003. Secondary Models. In Modeling Microbial Responses in Food, 1st ed. (ed. McKellar R, Lu X), pp. 63–150. CRC Press, BocaRaton.

Schmid AK, Pan M, Sharma K, Baliga NS. 2011. Two transcription factors are necessary for iron homeostasis in a salt-dwelling archaeon. Nucleic Acids Research, 39(7):2519–2533.

Schmid AK, Reiss DJ, Pan M, Koide T, Baliga NS. 2009. A single transcription factor regulates evolutionarily diverse but functionally linked metabolic pathways in response to nutrient availability. Molecular Systems Biology, 5:282. DOI: 10.1038/msb.2009.40.

Schnute J. 1981. A Versatile Growth Model with Statistically Stable Parameters. Canadian Journal of Fisheries and Aquatic Sciences, 38(9):1128–1140.

Sekse C, Bohlin J, Skjerve E, Vegarud GE. 2012. Growth comparison of several Escherichia coli strains exposed to various concentrations of lactoferrin using linear spline regression. Microbial Informatics and Experimentation, 2(1):5. DOI: 10.1186/2042-5783-2-5.

Sharma K, Gillum N, Boyd JL, Schmid A. 2012. The RosR transcription factor is required for gene expression dynamics in response to extreme oxidative stress in a hypersaline-adapted archaeon. BMC Genomics, 13:351. DOI: 10.1186/1471-2164-13-351.

Solak E, Murray-smith R, Leithead WE, Leith DJ, Rasmussen CE. 2003. Derivative Observations in Gaussian Process Models of Dynamic Systems. In Advances in Neural Information Processing Systems (ed. Becker S., Thrun S., Obermayer K.), volume 15, pp. 1057–1064. MIT Press.

Stephen DWS, Rivers SL, Jamieson DJ. 1995. The role of the YAP1 and YAP2 genes in the regulation of the adaptive oxidative stress responses of Saccharomyces cerevisiae. Molecular Microbiology, 16(3):415–423.

Tipping ME. 2001. Sparse Bayesian learning and the relevance vector machine. The Journal of Machine Learning Research, 1:211–244.

Todor H, Dulmage K, Gillum N, Bain JR, Muehlbauer MJ, Schmid AK. 2014. A transcription factor links growth rate and metabolism in the hypersaline adapted archaeon Halobacterium salinarum. Molecular Microbiology, 93(6):1172–1182.

Todor H, Gooding J, Ilkayeva OR, Schmid AK. 2015. Dynamic Metabolite Profiling in an Archaeon Connects Transcriptional Regulation to Metabolic Consequences. PloS One, 10(8):e0135693. DOI: 10.1371/journal.pone.0135693.

Todor H, Sharma K, Pittman AMC, Schmid AK. 2013. Protein-DNA binding dynamics predict transcriptional response to nutrients in archaea. Nucleic Acids Research, 41(18):8546–8558.

Tonner PD, Pittman AMC, Gulli JG, Sharma K, Schmid AK. 2015. A regulatory hierarchy controls the dynamic transcriptional response to extreme oxidative stress in archaea. PLoS Genetics, 11(1):e1004912. DOI: 10.1371/journal.pgen.1004912.

Verghese J, Abrams J, Wang Y, Morano KA. 2012. Biology of the Heat Shock Response and Protein Chaperones: Budding Yeast (Saccharomyces cerevisiae) as a Model System. Microbiology and Molecular Biology Reviews, 76(2):115–158.

Yao AI, Facciotti MT. 2011. Regulatory multidimensionality of gas vesicle biogenesis in Halobacterium salinarum NRC-1. Archaea, 2011:716456. DOI: 10.1155.2011.716456.

Yoon SH, Reiss DJ, Bare JC, Tenenbaum D, Pan M, Slagel J, Moritz RL, Lim S, Hackett M, Menon AL, et al. 2011. Parallel evolution of transcriptome architecture during genome reorganization. Genome Research, 21(11):1892–1904.

Yoon SH, Turkarslan S, Reiss DJ, Pan M, Burn JA, Costa KC, Lie TJ, Slagel J, Moritz RL, Hackett M, et al. 2013. A systems level predictive model for global gene regulation of methanogenesis in a hydrogenotrophic methanogen. Genome Research, 23(11):1839–1851.

You C, Okano H, Hui S, Zhang Z, Kim M, Gunderson CW, Wang Y, Lenz P, Yan D, Hwa T. 2013. Coordination of bacterial proteome with metabolism by cyclic AMP signalling. Nature, 500(7462):301–306.

Zhu Y, Kumar S, Menon AL, Scott RA, Adams MWW. 2013. Regulation of iron metabolism by Pyrococcus furiosus. Journal of Bacteriology, 195(10):2400–2407.

Zuber P. 2009. Management of oxidative stress in Bacillus. Annual Review of Microbiology, 63:575–597.

Zwietering MH, Jongenburger I, Rombouts FM, van’t Riet K. 1990. Modeling of the bacterial growth curve. Applied and Environmental Microbiology, 56(6):1875–1881.

